# Effects of calcium-regulated autophagy on *Candidatus* Liberibacter solanacearum in carrot psyllid midguts

**DOI:** 10.1101/2022.04.30.490068

**Authors:** Poulami Sarkar, Ola Jassar, Murad Ghanim

## Abstract

*Candidatus* Liberibacter solanacearum (CLso) transmitted by the carrot psyllid, *Bactericera trigonica* causes carrot yellows in Israel, and has recently gained much importance due to the excessive economical loss. Understanding the interactions between CLso and the psyllid at the cellular level is fundamental for the disease management. Here, we demonstrate the role of calcium ATPase, cytosolic calcium and most importantly Beclin1 in regulating autophagy and its association with Liberibacter. Presence of CLso generates reactive oxygen species and induces the expression of the detoxification enzymes in the psyllid midguts. CLso also induces the expression of both sarco/endoplasmic reticulum Ca^2+^ pump (SERCA) and 1,4,5-trisphosphate receptors (ITPR) in the midguts, followed by high levels of calcium in the cytosol. Silencing these proteins individually disrupted the calcium levels in the cytosol leading to direct effects on autophagy and thus on Liberibacter. On the other hand, inhibiting Beclin1-phosphorylation through different calcium induced kinases altered the expression of autophagy and CLso abundance. This study establishes a direct correlation between cytosolic calcium levels, autophagy and CLso in the carrot psyllid midgut.

## Introduction

Autophagy (also known as macroautophagy) is an evolutionary cellular process of recycling intercellular components and maintaining cellular functions. It plays a protective role during ER stress to protect the cells from metabolic damage and is essential for cellular homeostasis and development (1). It initiates with the formation of double-membrane vesicles known as autophagosomes (APs) from the ER (endoplasmic reticulum) involving vesicular engulfment of materials and pathogens or foreign particles, and fusion with lysosomes for lysosomal hydrolases and catabolic processes (2, 3). This in turn reduces ER stress consequently regulating apoptosis, and this coordination between autophagy and apoptosis is crucial for regulating cell survival or cell-death, respectively (4–6). High cytosolic calcium (Ca^2+^) during ER stress is one of the multiple signaling molecules regulating the induction of autophagy (7, 8). 1,4,5-trisphosphate receptors (ITPRs/IP3Rs) are tetrameric Ca^2+^ channels located at ER, which release Ca^2+^ from the ER to the cytosol and cytosolic Ca^2+^ re-enters the ER through a Ca^2+^ pump called ATP2A/SERCA (sarco/endoplasmic reticulum Ca^2+^) (9–13).

Cytosolic Ca^2+^ plays an important role as a pro-autophagic signal encompassing both Beclin1 and mTOR (mechanistic target of rapamycin) signalling cascades (14, 15). Beclin1, an ortholog of yeast Atg6, represents a determining link between autophagy and apoptosis, and is crucial for the initiation of autophagosome formation. Beclin1 when bound to Bcl2 inhibits autophagy and induces mitochondrial apoptosis. This interaction is dynamically regulated by death-associated protein kinase (DAPK), which phosphorylates Beclin1 at Thr119 antagonizing the interaction, thereby inducing autophagy (16–18). Serine/threonine kinase mTOR, in particular complex1 (mTORC1) is the master negative regulator of autophagy, which is inhibited by AMPK (AMP-activated protein kinase) which in turn is activated by Ca^2+^-calmodulin-dependent protein kinase kinase-β (CaMKKβ) upon increase in cytosolic Ca^2+^ (Ca2+-CAMKK2-AMPK pathway). AMPK also regulates autophagy by phosphorylating Beclin1 at Thr388 (18, 19). Active mTORC1 triggers inactivation of ULK (Unc51-like kinase, homolog of ATG1), a serine/threonine kinase that plays a key role in inducing autophagy (20–22). Loss of mTORC1 activity induces activated ULK to initiate autophagy by phosphorylating Beclin1 at Ser15 (18). Hence, Beclin1 accounts for the core molecular machinery involved in autophagy.

Bacterial pathogens often induce ER stress and free cytosolic Ca^2+^ in the host cells which in consequence induces autophagy as a defense mechanism to the cellular stress (23, 24).

However, many pathogenic bacteria can manipulate the autophagic pathway and replicate inside the autophagosomal compartment (23, 25, 26). Psyllid-Liberibacter relationship is one such system where the molecular mechanism of interaction and pathogenesis is obscure. One of the Liberibacter species, *Candidatus* Liberibacter solanacearum (CLso), Haplotype D is a known gram negative, unculturable bacterium transmitted by the carrot psyllid, *Bactericera trigonica* in a circulative and persistent manner. Recent transcriptomics revealed induced ER stress and autophagic genes in the psyllids in response to Liberibacter (27–30). Hijacking host immunity is crucial for bacterial survival and replication inside the host cells and autophagy seems important to reduce bacteria-induced cellular (ER) stress and maintain homeostasis. In recent studies, (CLso) is reported to induce autophagy in the psyllid host, *B. trigonica* (31) and in *Diaphorina citri* (30), and repress apoptosis in *B. cockerelli* (32).

In this study, we investigated the role of Ca^2+^ signaling and autophagy in CLso propagation inside the psyllids. In addition to this, we also studied the role of phosphorylated Beclin1 in autophagy initiation. We found elevated cytosolic Ca^2+^ and increase in autophagy-regulated genes in the psyllid midguts in response to CLso infection. Blocking the Ca^2+^ channels and the protein kinases in the calcium signaling cascade drastically disturbed both Beclin1 regulation leading to alterations in autophagy and CLso levels.

## Results

### CLso induces ROS generation in psyllid midguts

Psyllid midguts showed an increased expression of reactive oxygen species (ROS) (Fig. 1A-C) with enhanced expression in the cytochrome P450 (C450) and the detoxification enzyme, mainly superoxide-dismutase (SOD). However, there was a decline in the expression of glutathione S-transferase (GST) (Fig. 1B&C). Increased ROS was observed in CLso+ psyllid midgut nuclei (bright fluorescent red) upon DHE intercalation in response to higher oxidation (Fig. 1A).

**Fig. 1.**
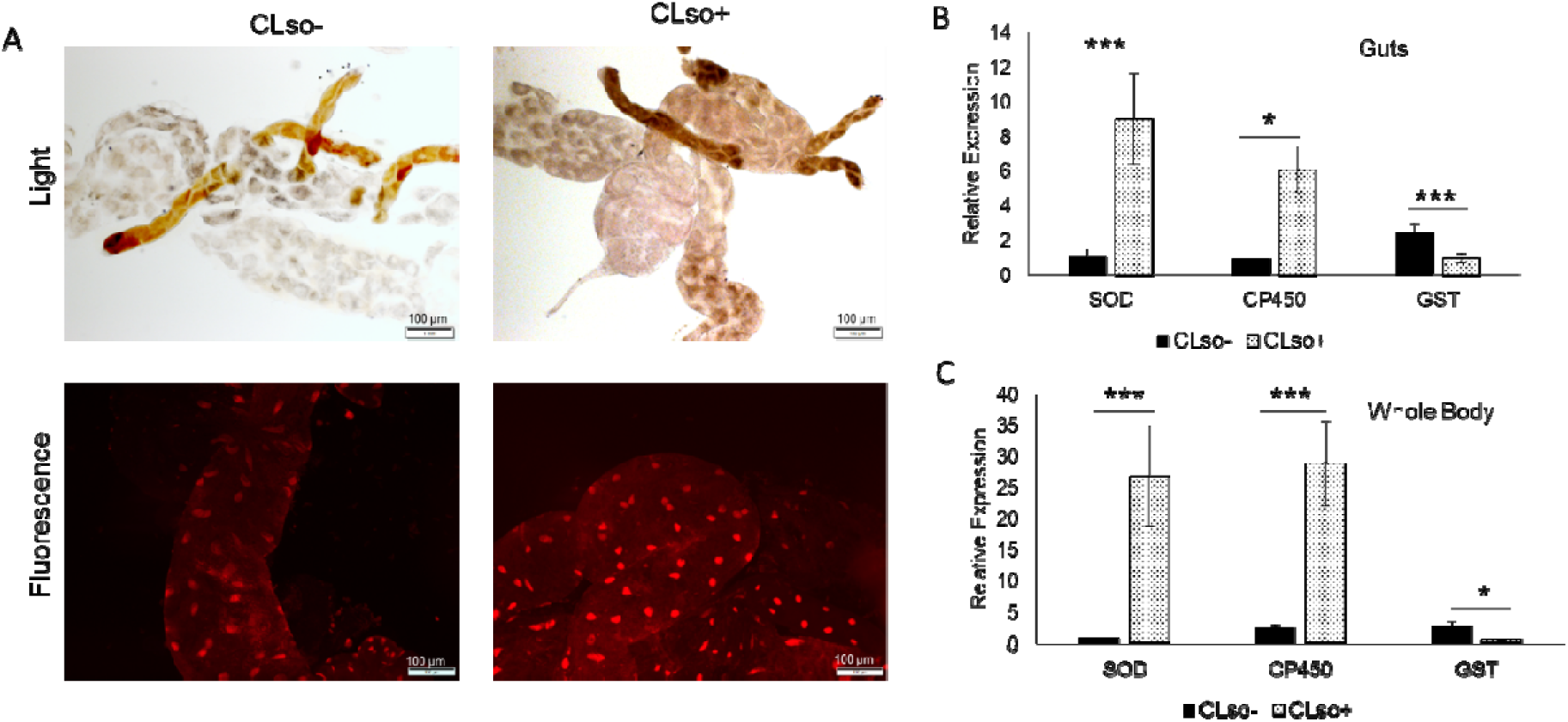
ROS detection in CLso-infected psyllid midguts. **A**. Light and fluorescence detection of ROS using DHE stain. **B**. Real-time analysis for SOD, CP450 and GST in guts and **C**. in whole psyllids. * indicates p ≤ 0.05 and *** indicates p ≤ 0.01.

### Induced expression of Calcium ATPases and calcium signaling genes with increased cytosolic calcium levels in CLso infected psyllids

*In situ* expression of SERCA (sarco/endoplasmic reticulum Ca^2+^) which is responsible for calcium influx from the cytosol to the ER, was observed using immunolocalization with SERCA antibody. Expression of SERCA was highly induced in CLso+ psyllid midguts as seen in Fig. 2A where it was observed around the nuclei stained with DAPI (blue), mimicking ER. Higher amounts of calcium was also detected in the CLso+ midgut cytosol when stained with calcium staining fluophore Fluo-8AM (Fig. 2B). Real-time expression analysis revealed upregulation of SERCA as well as some of the calcium induced genes in the calcium signalling pathway that leads to autophagy (Fig. 2B &C). The intensities of the signal verified the results (Fig. S1 A&B).

**Fig. 2.**
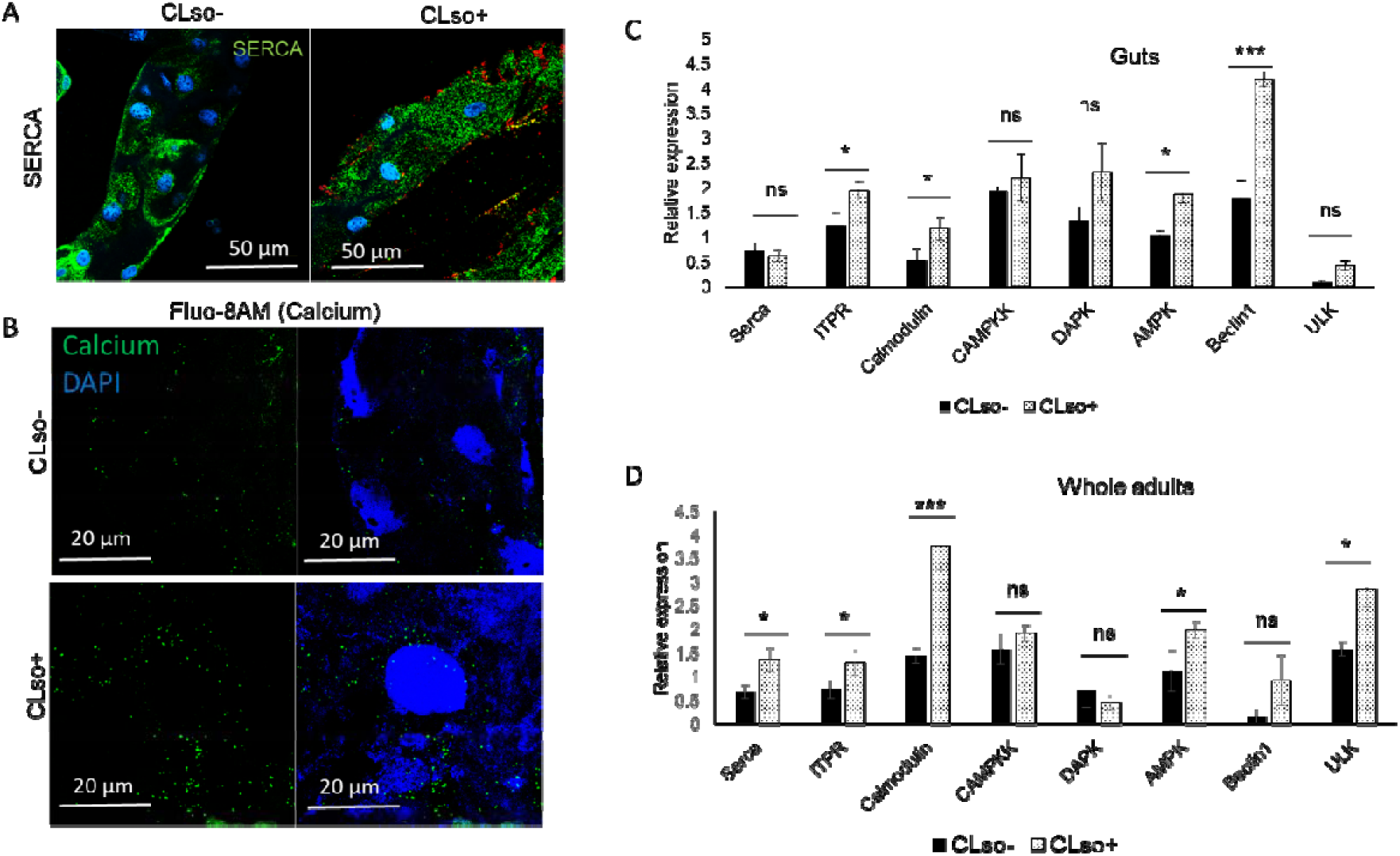
Differential expression of SERCA, calcium and calcium-signaling genes. **A**. Immunostaining of SERCA (green) and CLso (red) in CLso-free (CLso-) and CLso-infected (CLso+) psyllid midguts counterstained with DAPI (blue). **B**. Detection of cytosolic calcium levels (green) using Fluo-8AM staining in CLso- and CLso+ psyllid midguts, counterstained with DAPI (blue). **C**. Real-time analysis for the expression of SERCA, ITPR and calcium-signaling cascade genes in the midguts and **D**. in whole psyllids. * Indicates p ≤ 0.05, *** indicates p ≤ 0.01 and ns indicates not-significant.

### Inhibition of the calcium pumps in the ER membrane alters CLso titer

To study the role of calcium ATPases and calcium in CLso abundance, we silenced SERCA (responsible for calcium influx) and ITPR (responsible for calcium efflux) individually with double-stranded RNA (dsRNA). dsSERCA treated CLso+ psyllids showed reduction in SERCA gene expression in the midguts in both real-time and immunolocalization studies. (Fig. 3A&B). This resulted in increased accumulation of cytosolic calcium in the midguts as analysed using calcium binding fluophore, Fluo-8AM (Fig. 3C). Moreover, CLso titers drastically reduced as a result of silenced SERCA and consequent high calcium levels in the cytosol (Fig. 3A&B). The intensities of the signals were quantified using ImageJ as shown in Fig. S1C&D. Lysotracker was used to verify the increase in autolysosomes in the psyllid midguts (Fig. S2). This also resulted in decrease in apoptosis as seen with TMR-Red staining (Fig. 3D).

**Fig. 3.**
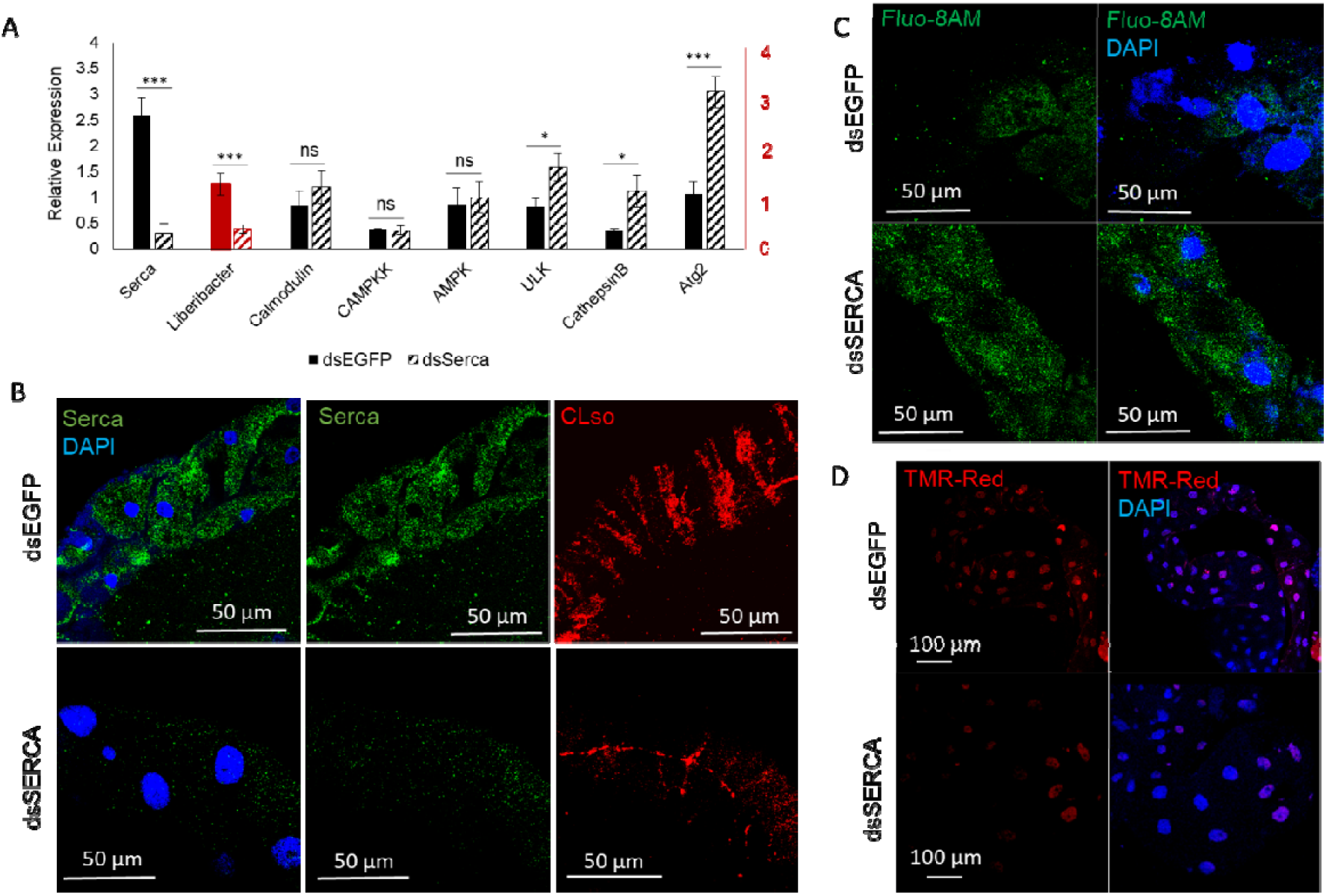
Effect of dsSERCA on CLso and autophagy. **A**. Real-time analysis of the differential expression change in SERCA and corresponding calcium-signaling genes along with CLso abundance in the psyllid midguts following dsSERCA treatment. * denotes p ≤ 0.05, *** indicates p ≤ 0.01 and ns indicates not-significant. **B**. Representative image of immunostaining analysis of SERCA (green) and CLso (red) in the midguts upon dsSERCA treatment, counterstained with DAPI (blue). **C**. Elevated levels of cytosolic calcium (green) in dsSERCA treated midguts. **D**. TUNEL assay showing reduced apoptosis in dsSERCA treated midguts.

On the contrast, dsITPR silenced midguts showed a reduction in cytoplasmic calcium levels and reduction in corresponding genes involved in the calcium signaling pathway along with increased CLso abundance in the midguts (Fig. 4A-C). The intensities of the signals for CLso were also analysed using ImageJ (Fig. S1E&F). The decrease in autophagy was also verified using Lysotracker staining (Fig.S2). Moreover, when the CLso+ psyllid midguts were stained for TMR-Red, there was an increase in apoptosis upon silencing ITPR expression (Fig. 4D).

**Fig. 4.**
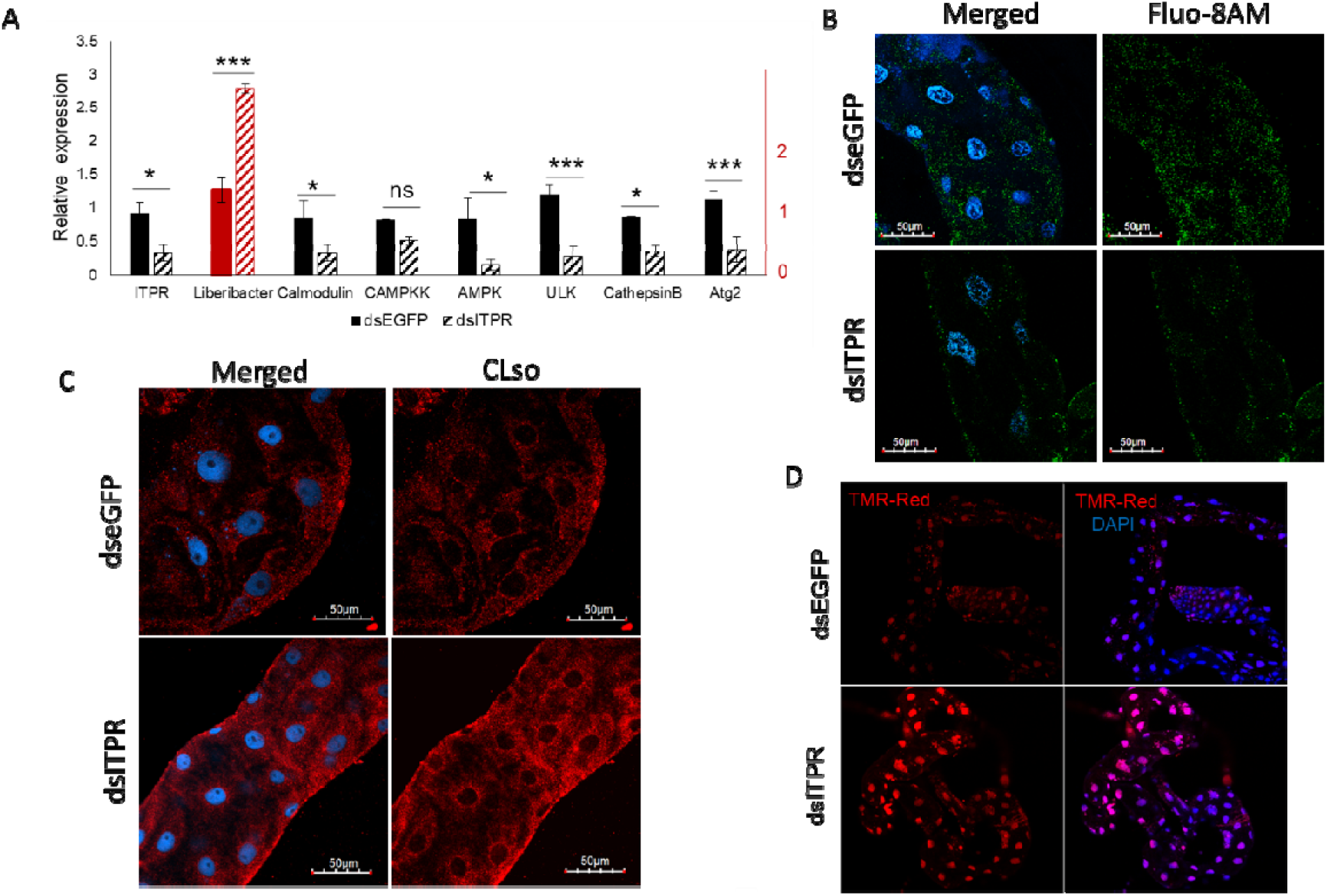
Effect of ITPR silencing on CLso and autophagy. **A**. Real-time analysis showing differential expression levels of ITPR, calcium-signaling genes and CLso abundance in the psyllid midguts following dsITPR treatment. * denotes p ≤ 0.05, *** denotes p ≤ 0.01 and ns indicates not-significant. **B**. Fluo-8AM staining of cytosolic calcium (green) after dsITPR treatment, counterstained with DAPI (blue). **C**. Immunostaining of CLso (red) in the psyllid midguts following dsITPR treatment. **D**. Increased apoptosis in dsITPR treated midguts as detected by TUNEL assay using TMR-red (red) and DAPI (blue).

In addition, Liberibacter was seen to be localized at the surface of the midgut cells in a stripe like pattern at one focal plane and around the nuclei as the focal plane is changed (Fig. S3).

### Inhibition of Beclin1 signaling cascade

Beclin1 phosphorylation was reduced *in vivo* by chemically inhibiting AMPK and DAPK which phosphorylates Beclin1 at sites Ser93,Ser96 and Thr119, respectively. Phosphorylation of both the sites Thr119 and Ser93,Ser96 was observed to be more in the CLso-infected psyllid midguts than in CLso-free psyllid midguts (Fig. 5&6). The intensity measurements of the signals were also calculated using ImageJ as shown in Fig. S1G&H. As expected, Beclin1 phosphorylation at sites Ser93, Ser96 was drastically reduced when AMPK was inhibited.

**Fig. 5.**
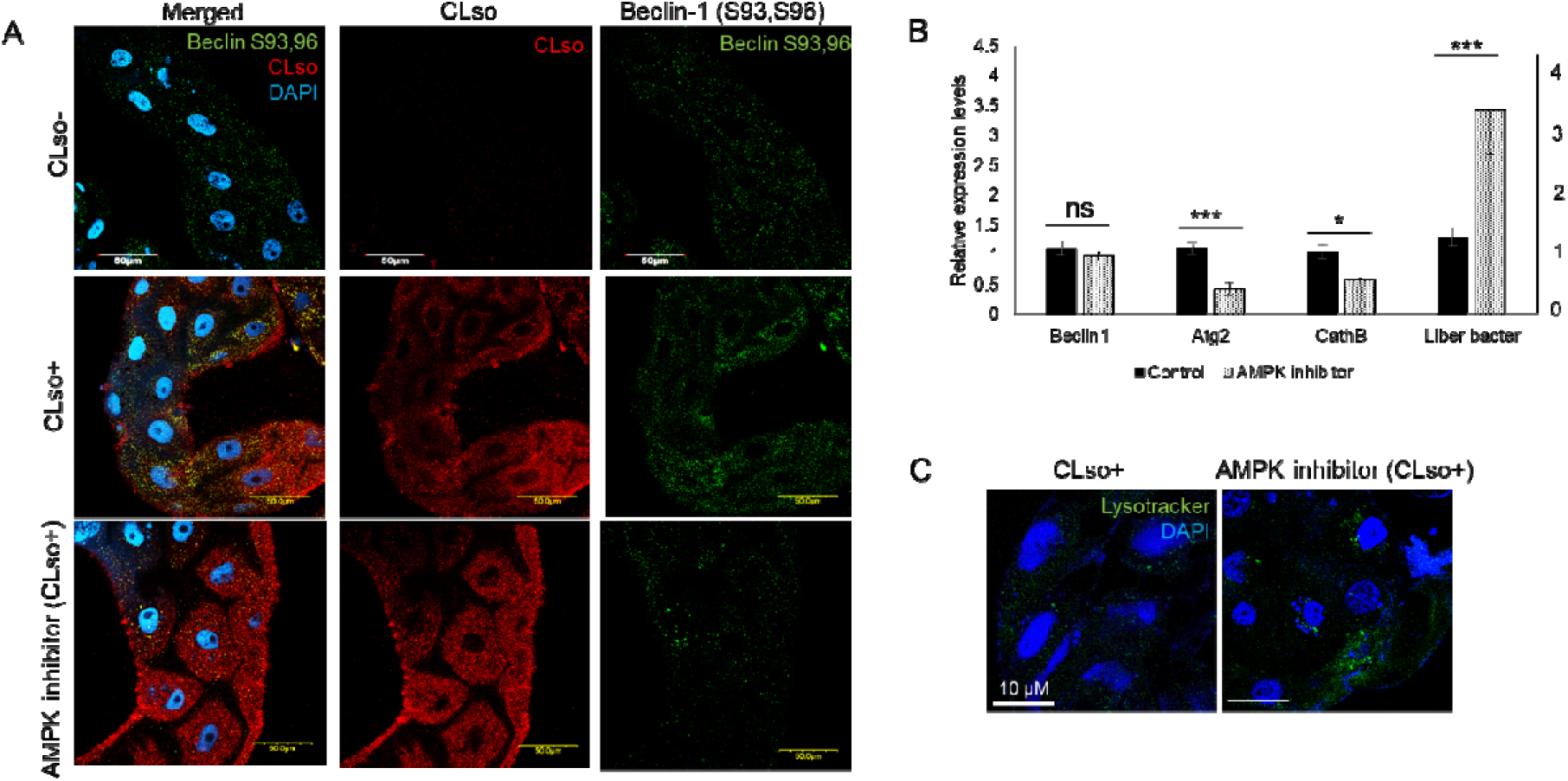
Effect of AMPK inhibitor on Beclin1 phosphorylation, autophagy and CLso. **A**. Immunostaining analysis reveals reduced Beclin1 phosphorylation (green) at Ser93,96 sites and higher accumulation of CLso (red) in the AMPK inhibited psyllid midguts, counterstained with DAPI (blue). **B**. Real-time assay for the differential expression of Beclin1, autophagy genes and CLso abundance following AMPK inhibition. p ≤ 0.05 is indicated by *, p ≤ 0.01 by *** and ns indicates not-significant. **C**. Autolysosome detection using Lysotracker DND (green) counterstained with DAPI (blue).

**Fig. 6.**
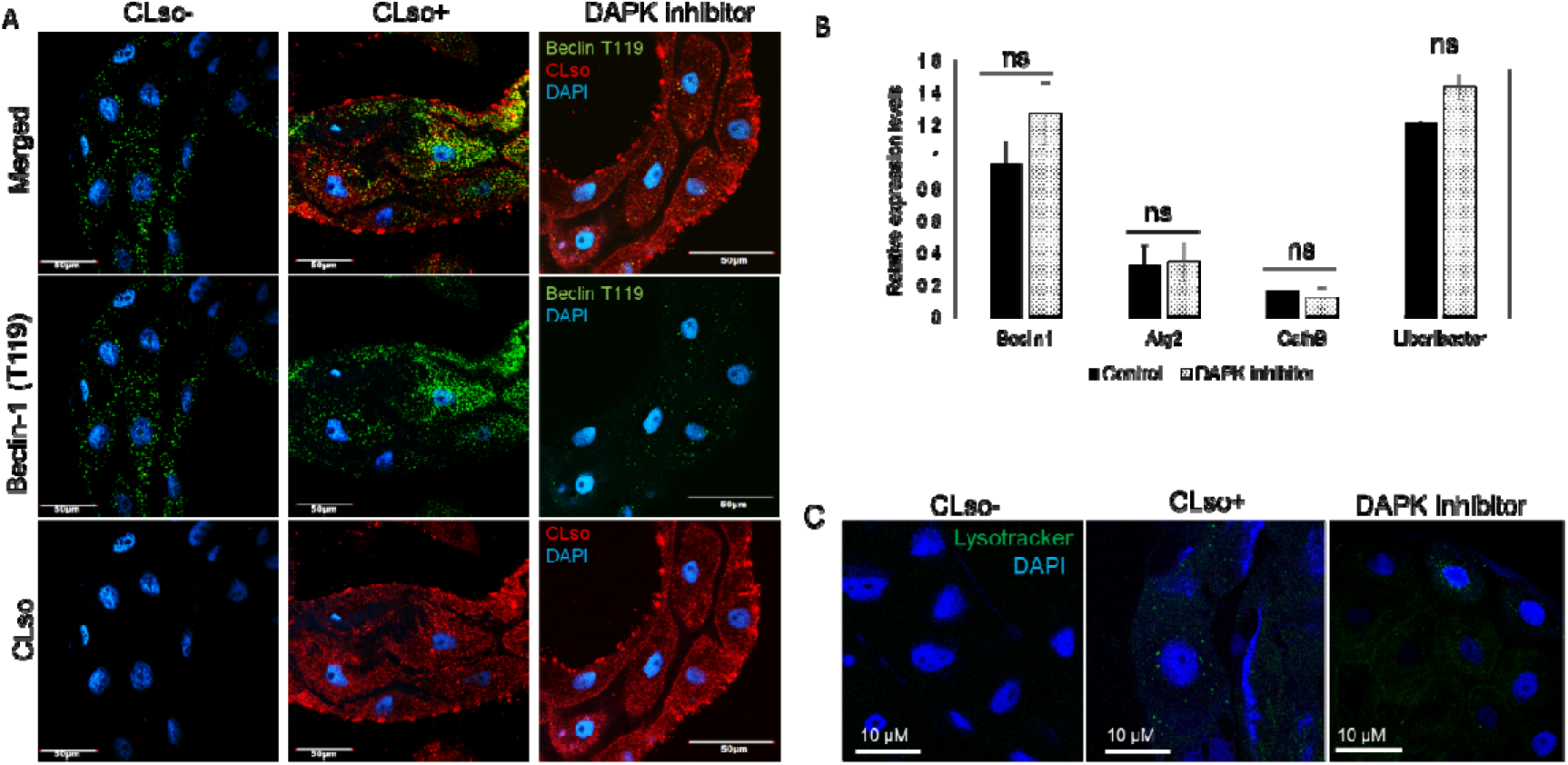
Effect of DAPK inhibitor on Beclin phosphorylation, autophagy and CLso abundance. A. Immunostaining analysis for Beclin1 phosphorylation at site Thr119 (green) and CLso (red) counterstained with DAPI (blue). **B**. Real-time analysis showing differential expression levels for Beclin1, autophagy genes and CLso titre. ns denotes not-significant. **C**. Autolysosome detection using Lysotracker DND (green) counterstained with DAPI (blue).

This consequently resulted in increase in CLso abundance in the midguts. We also checked the effect of this inhibition on autophagy, and the autophagy-related genes, specifically Atg2 was downregulated (Fig. 5). Additionally, the presence of lysosomes was reduced which indicated less autophagy (Fig. 5C).

On the contrast, for DAPK inhibition, there were no significant changes in CLso abundance even when Beclin1 remained un-phosphorylated at site Thr119 as seen in Fig 6A&B. Autophagy-related genes and LysoTracker staining confirmed no significant change in the midguts following DAPK inhibition (Fig. 6B&C).

## Discussion

*Candidatus* Liberibacter solanacearum (CLso), Haplotype D is an intracellular, gram-negative bacteria transmitted by the carrot psyllid, *Bactericera trigonica*. Understanding the stress response and innate immunity of the vector is crucial to determine new approaches to disrupt the disease transmission. Innate immunity involves both autophagy and apoptosis and functions to maintain homeostasis during pathogenesis (4, 33, 34). Autophagy maintains cellular homeostasis during ER stress by eliminating intracellular pathogens and delivering it to lysosomes for destruction (2, 3, 35, 36). One of the major signaling molecules regulating autophagy is calcium (Ca2+) ions which signals Beclin1 through different protein kinases present in the calcium signaling cascade that leads to autophagy initiation (7, 8, 14, 15). A number of recent studies involving Liberibacter species have demonstrated the occurrence of autophagy in the insect vector during infection (30, 31, 37). CLas induces the formation of a double-membrane autophagic vacuoles from the ER membrane (30). In addition, major autophagy-related genes are reported to be upregulated in Asian citrus psyllid (30) and the potato psyllid (37) following Liberibacter infection.

In this study, we tried to explore the role of calcium and the calcium signaling cascade proteins involved in autophagy in carrot psyllids during CLso infection. As pathogenesis induces ROS, which in turn stimulates autophagy (38), we also analysed ROS generation in the psyllid midgut. DHE staining of the midguts show higher ROS generation in CLso-infected psyllids than in CLso-free psyllids (Fig. 1A). Real-time expression studies of detoxification enzymes show upregulation of superoxide dismutase and cytochrome P450 genes in CLso-infected psyllids with downregulation of glutathione S-transferase (Fig. 1B&C). Next, we checked the expression of Ca^2+^ ATPases and the cytosolic Ca^2+^ levels in the psyllid midguts upon CLso infection. Immunostaining of SERCA revealed higher intensity of signal in CLso-infected midguts (Fig. 2A) which also showed higher signals for cytosolic calcium (Fig. 2B). Expression of Ca^2+^ influx (SERCA) pumps were elevated in CLso-infected midguts along with other protein kinases involved in the calcium signaling pathway including Beclin1 (Fig. 2C&D). This elevation in SERCA could be in order to maintain calcium homeostasis for reducing ER stress caused by CLso. However, it’s interesting to see high levels of cytosolic calcium despite overexpression of SERCA. We checked the expression of Ca^2+^ efflux pump protein, ITPR which was also seen to be overexpressed in the CLso-infected psyllids (Fig. 2C&D).

Next, to understand the involvement of these two pumps in CLso propagation and autophagy, we silenced both SERCA and ITPR individually using double-stranded RNA. Silencing SERCA was validated using both real-time expression analysis and immunostaining (Fig. 3A&B). Midguts treated with dsSERCA showed an increased levels of cytosolic calcium (Fig. 3C) as well as elevated levels of calcium signaling genes, autophagy-related gene-2 and Cathepsin-B (Fig. 3A). This also resulted in high number of autolysosomes (Fig. S2) and reduction in apoptosis (Fig. 3D). As it is known that autophagy and apoptosis cross-regulate each other (4–6), our results verified upregulation in autophagy upon elevated cytosolic Ca^2+^ levels caused by silencing SERCA. Consequently, there was a reduction in the levels of CLso in dsSERCA treated midguts (Fig. 3B). Comparably, when ITPR was silenced using dsRNA, the calcium signaling genes and autophagy-related gene2 were downregulated (Fig. 4A) along with reduced cytosolic Ca^2+^ (Fig. 4B). This resulted in increased levels of CLso in the dsITPR treated midguts (Fig. 4C). Corresponding LysoTracker and TUNEL assay confirmed higher apoptosis (Fig. 4D) and reduction in autophagy (Fig. S2) in the dsITPR treated midguts.

As Beclin1 is crucial for autophagy and leads to initiation of autophagosome formation, we wanted to examine the role of two important protein kinases involved in the calcium signaling pathway in Beclin1-mediated autophagy. As depicted in Fig. 7, cytosolic Ca^2+^ activates both death-associated protein kinase (DAPK) and AMP-activated protein kinase (AMPK), which in turn phosphorylates Beclin1 at specific sites to initiate autophagosome formation. In this study, we tried to chemically inactivate both DAPK and AMPK individually, to see which protein kinase is crucial for Beclin1-mediated autophagy. AMPK inhibitor drastically reduced the phosphorylation of Beclin1 at S93,96 sites, leading to accumulation of CLso in the treated midguts (Fig. 5A). This also led to reduction in autophagy as analysed using real-time analysis (Fig. 5B) and Lysotracker (Fig. 5C). On the other hand, DAPK inhibitor reduced the phosphorylation of Beclin1 at site Thr119 (Fig. 6A), however, there was no significant change in CLso abundance (Fig. 6A&B) or in autophagy-related genes (Fig. 6B&C).

**Fig. 7.**
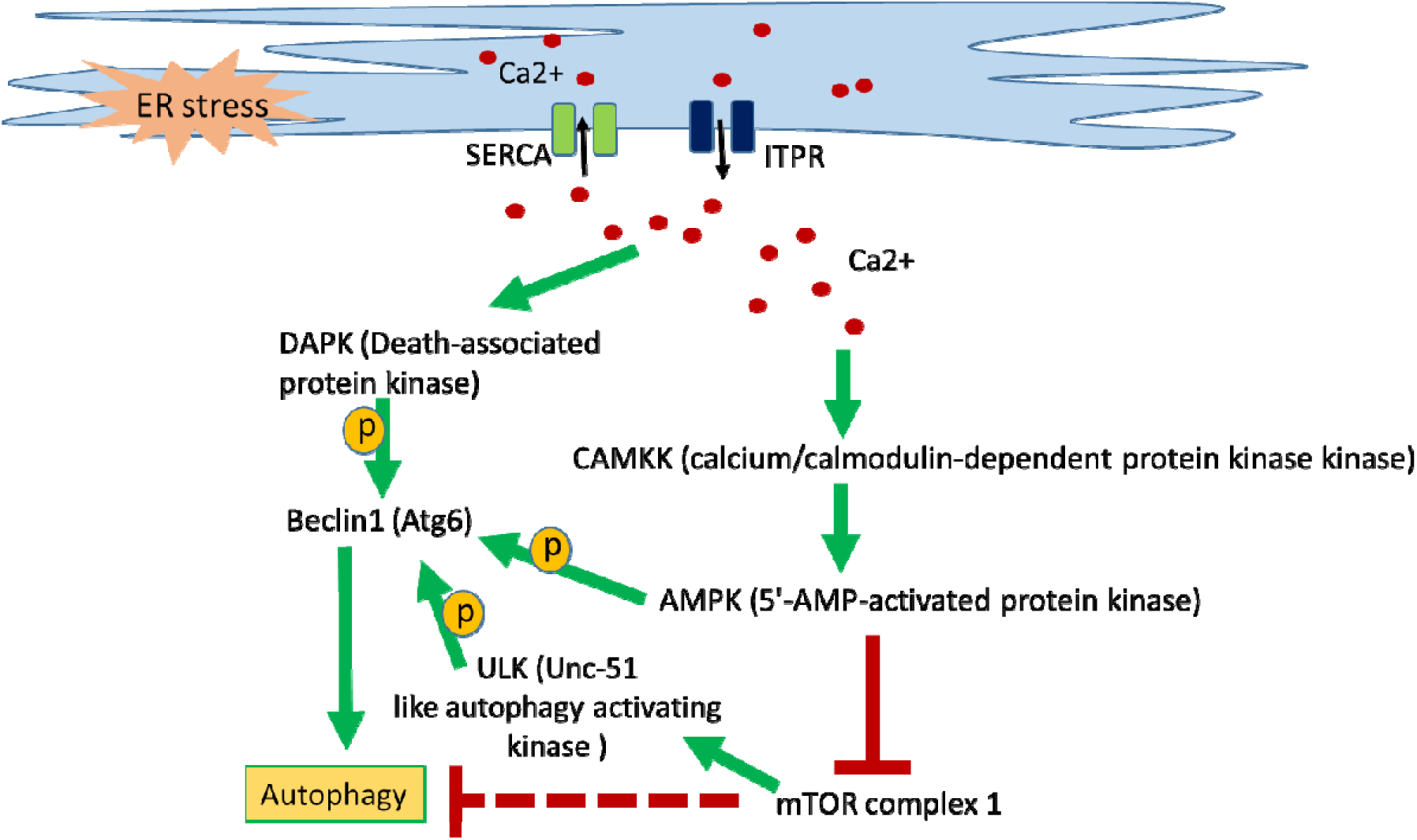
Depiction of calcium signalling cascade leading to autophagy. Calcium influx (SERCA) pumps and efflux (ITPR) pumps maintain the calcium homeostasis in the ER as well as in the cytosol. On the other hand, ER stress causes elevated cytosolic calcium levels, which in turn activates downstream protein kinases leading to activation of Beclin1 which helps in the initiation of autophagophore formation leading to autophagy.

This study demonstrated that cytosolic Ca^2+^ regulates autophagy through activating AMPK. It also revealed that AMPK is crucial for Beclin1 functioning in autophagy initiation and that autophagy and CLso propagation negatively regulate each other to maintain a survival balance of both the pathogen and the vector. It will be further interesting to explore the exact mechanism leading to elevated calcium levels in the cytosol and to know whether autophagy is a response to Liberibacter as a part of innate immunity.

## Methods and materials

### Insects and plants material used

*Ca*. Liberibacter solanacearum - infected (CLso+) and *Ca*. Liberibacter solanacearum - free (CLso-) psyllids were maintained on parsley (*Petroselinum crispum*) in separate cages under 14h photoperiodic light, 25±2 °C and 60 % humidity. Psyllids were periodically tested for CLso using PCR analyses.

### ROS imaging

In situ ROS detection was carried out using dihydroethidium (DHE) (Sigma-Aldrich, Israel). Briefly, midguts were dissected out of psyllids in PBS (phosphate buffer saline), and immediately incubated in 10mM DHE for 7 minutes in the dark. The midguts were washed with PBS thrice before mounting in glass slides and viewed under confocal microscope.

### dsRNA preparation and treatment

dsRNA for SERCA (dsSERCA) and ITPR (dsITPR) and control dsRNA (ds-eGFP) was prepared using the T7 FlashScribe transcription kit (Cellscript, USA). PCR amplified products using gene-specific primers containing T7 promoters on 5′ end was used for dsRNA production (Table1). dsRNA quality and quantity was checked using agarose gel electrophoresis and NanoDrop 1000 spectrophotometer (Thermoscientific). dsRNA feeding was carried out using fresh leaf flush as previously described (39). ds-eGFP was used as a control. Each experiment was replicated thrice with minimum of 10 insects for each treatment.

**Table 1.**
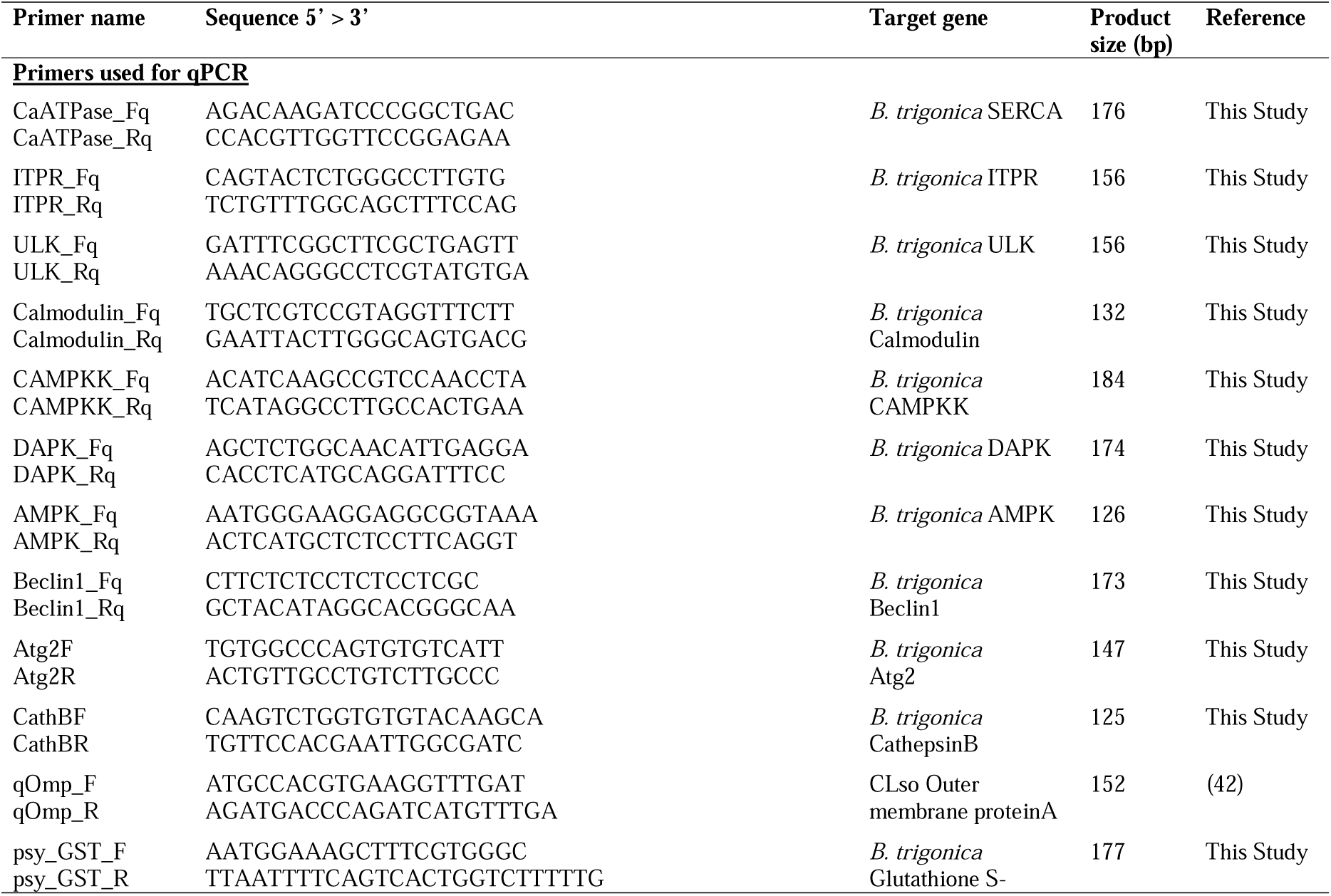
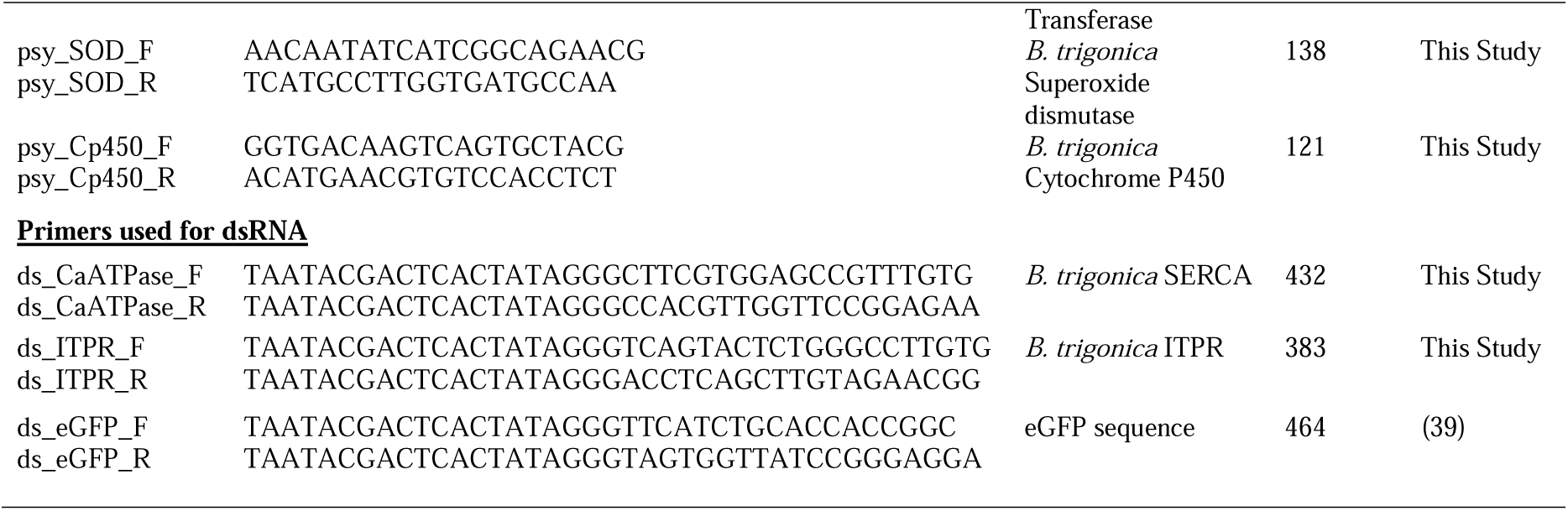

### DAPK and AMPK inhibition

To identify which pathway is most important for Beclin1 mediated autophagy, Beclin1-phosphorylation by DAPK and AMPK was inhibited using DAPK inhibitor and AMPK inhibitor (Merck), respectively. 10 ug/ml of inhibitor in ethanol was applied on a fresh parsley flush with around 20 psyllids in a jar. After 16h the insects were collected for DNA/RNA isolation or dissection for microscopy. Ethanol was applied to a flush as a control experiment. Each experiment was replicated thrice with minimum of 15 insects for each treatment. Beclin1 phosphorylation corresponding to each treatment was detected using antibodies as described later in this study.

### Calcium staining and LysoTracker staining

To detect the cytosolic calcium, we used Fluo-8AM (Abcam). Briefly, the midguts were dissected in PBS and incubated with Fluo-8AM in HBSS (Hank’s balanced salt solution) for 30 min for 1hr in 37 °C in dark. The midguts were then washed with PBS thrice and was mounted with DAPI for microscopy. Auto-lysosomes were detected using LysoTracker Green DND-26 (Invitrogen) as described previously (31).

### Terminal Deoxynucleotidyl Transferase dUTP Nick End Labelling (TUNEL-assay)

The psyllid midguts were diagnosed for cell-death or apoptosis using *In situ* cell-detection kit TMR-red (Roche) following the manufacturer’s instructions. Briefly, the midguts were fixed in 4 % paraformaldehyde in PBS for 1 h followed by incubation in permeabilization solution (0.1 % Triton X-100) for 15 min. 100 µl of Label solution was used as a negative control. 50 µl of Enzyme solution was mixed with 450 µl of Label solution to form a TUNEL reaction mixture in which the midguts were incubated for 1 h at 37 °C in the dark followed by washing thrice in PBS and finally the midguts were mounted in glass slides with DAPI to view under the microscope.

### Immunostaining analysis

Immunostaining for psyllid proteins as well as for liberibacter was done as described previously (39). Briefly, the midguts were dissected out in PBS, fixed in 4% paraformaldehyde, treated with Triton-X for 30min and blocked with 1.5 % BSA (bovine serum albumin) for 1 h. The midguts were then incubated with primary antibody for liberibacter with anti-OmpB-antibody (GenScript), followed by secondary antibody conjugated with Cy3/Cy5 (Jackson ImmunoResearch Laboratories). This was followed by incubation with psyllid protein antibodies for Beclin1, anti-phospho Beclin1-Ser93/Ser96 (Cell signaling Technologies) for AMPK inhibition experiment or anti-phospho Beclin1-Thr119 (Sigma Aldrich, Israel) for DAPK inhibition experiment and a secondary antibody conjugated to Cy3/Cy5. The midguts were washed at least thrice before mounting on a slide with DAPI and were visualized with Olympus IX81 confocal microscope.

### RNA isolation and qRT-PCR for gene expression

Total RNA was isolated from psyllid midguts and whole body from different experiments using Tri Reagent (Sigma Aldrich, Israel) as described previously. DNA contaminations were removed using DNaseI (Thermo Scientific). A minimum of 15 psyllids/ midguts were used for each experiment. First strand synthesis was carried out using M-MLV reverse transcriptase (Promega Corporations) following the manufacturer’s instructions. Real-time PCR was carried out using ABsolute Blue SYBR green mix in a StepOne real-time PCR system (Applied Biosystems). Ct values were normalized using Elongation factor 1α (Ef1α). The expression of each gene was calculated following Livak (2^-ΔΔCt^) method (40) for relative gene expression.

### Quantification of CLso and qPCR

To quantify the relative abundance of *Ca*. Liberibacter solanacearum from each treatment, total DNA was isolated from individual psyllid/ midgut from each experiment using modified CTAB protocol (41) as described in a previous study (39). Real-time analysis was carried out as described before using actin as a housekeeping gene. Each psyllid/ midgut was crushed in 250 µl/ 100 µl of CTAB buffer, respectively and were incubated for 1h at 37 °C followed by phenol-chloroform purification.

### Statistical analysis

The significance of relative expression analyses performed for both qRT-PCR and qPCR were determined using at least 10-12 samples each using one-way ANOVA with Tukey’s post hoc test (p<0.05).

## Notes

### Competing Interest Statement

The authors have declared no competing interest.

