## Supplementary material for "Effects of calcium-regulated autophagy on *Candidatus* Liberibacter solanacearum in carrot psyllid midguts": Fig. S

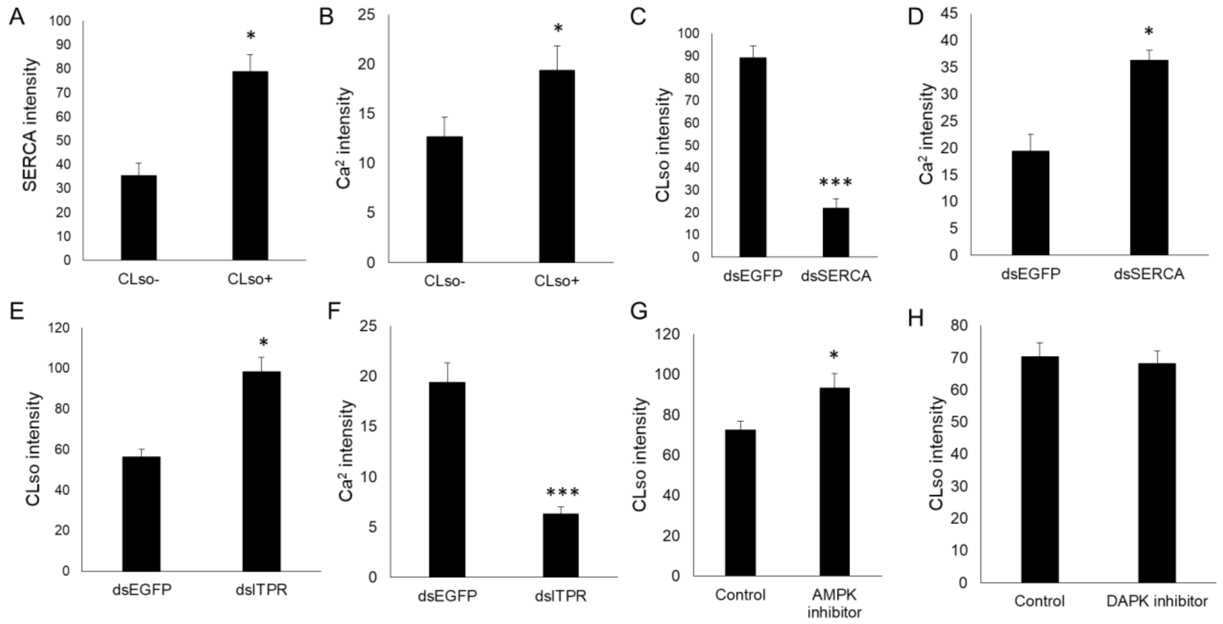


**Fig. S1** Intensity measurements for the confocal images using ImageJ. **A.** Immunostaining signals for SERCA antibody. **B.** Calcium signals for Fluo-8AM. **C.** Immunostaining signals for CLso following dsSERCA treatment. **D.** Calcium signals following dsSERCA treatment. **E.** Immunostaining signls for CLso after dsITPR treatment. **F.** Calcium signals following dsITPR treatment. **G.** Immunostaining signals for CLso following AMPK inhibitor. **H.** Immunostaining signals for CLso after DAPK inhibitor. * denotes $p \leq 0.05$, *** denotes $p \leq 0.0$; $n \geq 7$.

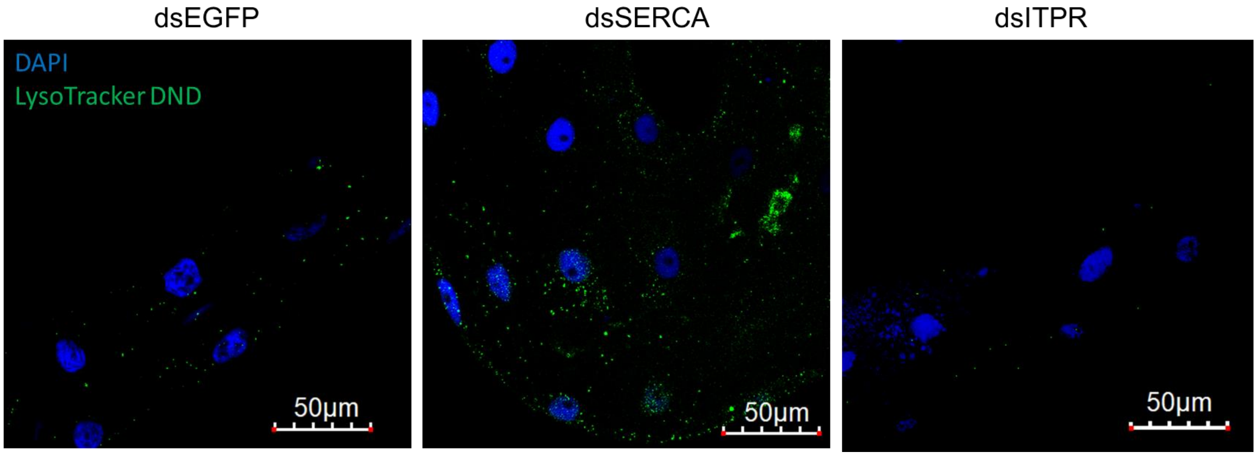


**Fig. S2** Detection of autolysosomes with Lysotracker DND (green) in dsSERCA and dsITPR treated midguts in comparision with dsEGFP treated midguts, counterstained with DAPI (blue).

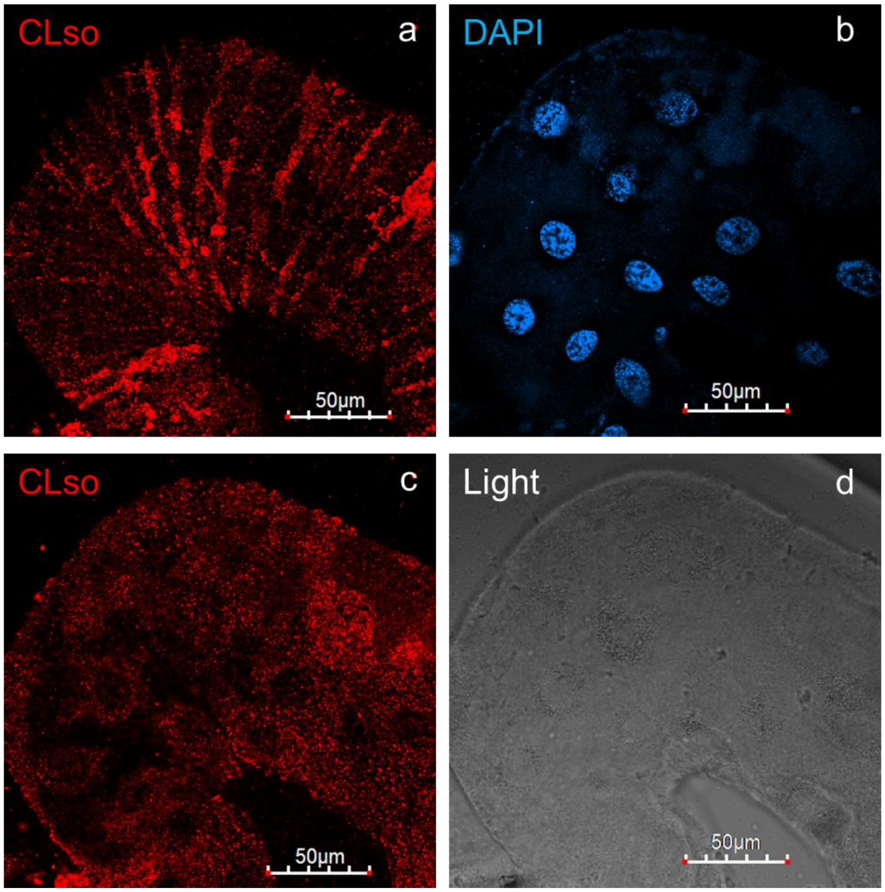


**Fig. S3** Localization of CLso at different focal planes as visualized in the midguts. CLso (red) localizes as stripes at the surface and around the nuclei (blue) of the midgut cells.
